# Identification of SAMD4B as a Centrosome-Associated Regulator of the Sonic Hedgehog Signaling Pathway

**DOI:** 10.64898/2026.11.19.767524

**Authors:** Ondřej Bernatík, David Vysloužil, Petra Paclíková, Tereza Renzová, Petra Kovaříková, Lukáš Čajanek

## Abstract

**Background:** Many protein-protein interactions are required for the assembly and function of primary cilia, microtubule-based organelles critical for the Sonic hedgehog (Shh) signaling pathway and, in turn, for embryonic development and tissue homeostasis. However, our understanding of this complex process remains incomplete.

**Methods:** To identify new regulators of primary cilia formation or function, we carried out proximity labeling experiments and compared interactomes under ciliogenesis-restrictive and permissive conditions. Using an unbiased strategy, we selected candidates that were tested functionally by siRNA-mediated depletion or gene editing, in combination with gene expression analysis and various microscopy techniques, using either vertebrate cell lines or zebrafish as model systems.

**Results:** We demonstrate that one of our identified candidates, protein SAMD4B, is associated with the centrosome and is required for Shh signaling in NIH3T3 cells and eye development in zebrafish. Our data implicate defective ciliary recruitment of Shh pathway regulators ARL13B and SMO as a plausible mechanism underlying the SAMD4B-mediated regulation of Shh signaling.

**Conclusions:** Together, our data identify a previously unrecognized link between SAMD4B, a protein known to be associated with cytoplasmic RNA condensates, and ciliary signaling.

## Background

The primary cilium is an antenna-like organelle found on the surface of most vertebrate cells. It plays important signaling roles that govern many key events during embryogenesis and contributes to tissue homeostasis in adulthood (Čajánek et al., 2025; Gopalakrishnan et al., 2023; Hilgendorf et al., 2024; Ott and Lippincott-Schwartz, 2025). In turn, defects in ciliary structure or function are associated with a heterogeneous group of disorders collectively termed ciliopathies (Mill et al., 2023; Reiter and Leroux, 2017).

The cilium comprises the basal body, transition zone, and microtubule-based axoneme enclosed by the ciliary membrane (Satir et al., 2010). The basal body originates from the mother centriole, the older of the two centrioles that form the centrosome. Unlike the daughter centriole, the mother centriole possesses distal appendages (composed of proteins such as CEP83/CCDC41, CEP89/CEP123, and CEP164) and subdistal appendages, specialized structures that are essential for ciliogenesis and microtubule anchoring, respectively (Bornens, 2012; Breslow and Holland, 2019). Some components of the centriole distal end, such as CP110, are displaced through distal appendage-dependent mechanisms to trigger axoneme formation (Spektor et al., 2007; Tanos et al., 2013); others, such as CEP104, subsequently localize to and define the tip of the nascent cilium (Satish Tammana et al., 2013).

The ciliary signaling function is supported by both the compartmentalization of the cilium from the rest of the cell and the dynamic trafficking of signaling components into and out of the organelle, allowing its molecular composition to be adapted to the requirements of specific signaling pathways (Hilgendorf et al., 2024; Mill et al., 2023; Nachury and Mick, 2019). The compartmentalization is ensured by the transition zone, a boundary between the basal body and the ciliary axoneme. Acting as a diffusion barrier and selective gate at the ciliary base, the transition zone regulates the entry of both membrane-bound and soluble proteins to maintain the distinct microenvironment of the cilium (Garcia-Gonzalo and Reiter, 2017; Park and Leroux, 2022). In addition, Intraflagellar transport (IFT) mediates bidirectional cargo trafficking between the ciliary base and tip. This process is driven by complexes of individual IFT proteins (such as IFT88, IFT81, IFT74, etc.) in cooperation with kinesin and dynein motors and is essential for cilium assembly, maintenance, and signaling (Lacey and Pigino, 2025; Prevo et al., 2017).

The Sonic hedgehog (Shh) signaling represents the prototypical pathway critically dependent on primary cilia (Bangs and Anderson, 2017; Huangfu and Anderson, 2005; Ingham, 2022). In the absence of Shh ligand, the Smoothened (SMO) receptor is present in the ciliary membrane at very low levels owing to repression by the Patched (PTCH) receptor, which restricts SMO access to its activating sterol ligands. As a result, SMO is ubiquitinated and removed from the cilium. Under these conditions, protein kinase A (PKA) phosphorylates GLI transcription factors, promoting their partial proteolytic processing into transcriptional repressor forms that suppress Shh target gene expression. Notably, proper localization dynamics of Shh signaling components inside cilia requires the activity of the small GTPase ARL13B (Caspary et al., 2007). Shh ligand binding to PTCH leads to removal of PTCH from the primary cilium, allowing SMO to accumulate within the ciliary membrane and gain access to activating sterols. In turn, activated SMO results in decreased PKA activity in the cilium. Consequently, GLI processing into repressor forms is blocked, enabling the activator forms to accumulate and induce transcription of Shh-responsive genes in the nucleus, including some of the pathway components such as *GLI* or *PTCH* (Hilgendorf et al., 2024; Ingham, 2022; Rohatgi et al., 2007). Proper activation of the Shh pathway is essential for neural tube patterning and for skeletal and eye development (Bangs and Anderson, 2017; Ingham, 2022).

Current estimates suggest that thousands of protein-protein interactions are required for cilium assembly and function (Arslanhan et al., 2020; Čajánek et al., 2025). However, our understanding of this complex process, including the identity of individual interactors, their specific roles, and their upstream regulation, remains limited. Moreover, an emerging theme in the field is the growing number of reports describing RNA-binding proteins and their regulators found in the vicinity of cilia and the centrosome, raising questions about their functions in these compartments (Gupta et al., 2015; Pachinger et al., 2025; Youn et al., 2018; Zhang et al., 2025). Here, using proximity-based proteomics followed by validation experiments, we identify SAMD4B, an RNA-binding protein found earlier in RNA-containing cytoplasmic condensates (Luo et al., 2010; Thomas et al., 2025; Wang and Zhang, 2023), as a centrosome-associated factor required for proper Shh signaling in NIH 3T3 cells and eye growth in zebrafish. To the best of our knowledge, this study provides the first evidence linking SAMD4B to the regulation of ciliary signaling.

## Materials and Methods

### Cell culture, transfections, and treatments

All cell lines were cultivated at 5% CO_2_ at 37°C. hTERT-RPE1, hTERT-RPE1 Flp-In T-Rex (Binó et al., 2022), and TTBK2 KO RPE-1 Flp-In T-Rex cells (Bernatik et al., 2020), were grown in DMEM F12 (ThermoFisher, Cat. No. 11320033) supplemented with 10 % FBS (BioSera, Cat. No. FB-1101/500). Tetracycline-free FBS (Biosera, Cat. No. FB-1001T/500) was used when inducible expression of the transgene by doxycycline was performed. The media were further supplemented with 1% Penicillin/Streptomycin (BioSera, Cat. No. XC-A4122/100) and 1% L-glutamine (BioSera, Cat. No. XC-T1715/100.). NIH 3T3 cells were cultured in GlutaMax DMEM (Thermo Fisher, Cat. No. 31966047) supplemented with 10 % FBS, and 1% Penicillin/Streptomycin. Normal human dermal fibroblasts (nHDF; Thermo Fisher, Cat. No. C0045C) were cultivated in KnockOut DMEM (Thermo Fisher, Cat. No.10829018) supplemented with 10% FBS, 1% L-glutamine, 1% nonessential amino acids (BioSera, Cat. No. XC-E1154), 1% Penicillin/Streptomycin, and 1% β-mercaptoethanol (Merck, Cat. No. M3148). To induce primary cilium formation, cells were serum-starved in medium containing 0-0.5% serum for 24 hours. Where indicated, SAG (Smoothened agonist; Merck, Cat. No. SML1314, or Tocris, Cat. No. 4366) was added in parallel during serum starvation to activate the Shh pathway (200 nM for NIH 3T3 cells and 500 nM for nHDF cells).

Plasmid transfection was carried out by Lipofectamine 3000 (Thermo Fisher, Cat. No. L3000008) according to the manufacturer’s instructions (up to 0.5 µg DNA per condition/well, 24-well plate format). To knock down mRNA expression, cell were transfected at the time of seeding (80 000 cells/well in a 24-well plate) using Silencer Select siRNA (mix of two siRNAs in concentration of 25nM each, or single siRNA in 40 nM concentration; Thermo Fisher) and Lipofectamine RNAiMAX (Thermo Fisher, Cat. No. 13778150) and OptiMEM (Thermo Fisher, Cat. No. 31985070) 72 hours before analysis. Fresh medium was exchanged on day 1. On day 2, the cells were treated and serum-starved as indicated, and in case of transfection with individual Samd4B siRNA #1, #2, or #3 transfected again with 15 nM of each siRNA. Used siRNAs are listed in Suppl. Table 3. siRNA against firefly luciferase (GL2) was used as a non-targeting control.

### Cloning

Human cDNA for CEP89 (Sillibourne et al., 2013), IFT81 (Bhogaraju et al., 2013), CP110 (Glatter et al., 2009), and CEP104 (gift from Vladimir Varga) was cloned into pENTR-TOPO vector using TOPO Gateway cloning kit (ThermoFisher, Cat. No. K450001). The obtained Gateway entry clones were verified by DNA sequencing, and the inserts were subsequently cloned into the pgLAP2-Neo-Myc-BirA (BioID) Gateway destination vector (Pejskova et al., 2020), using LR cloning kit (ThermoFisher, Cat.No. 11791020). Lentiviral vector for TTBK2 expression was custom cloned at Vectorbuilder, ID: VB190408-1222bfd (Benk Vysloužil et al., 2025).

### Stable cell lines

Generation of the stable cell lines of hTERT-RPE1 Flp-In™ T-Rex™ WT and TTBK2 KO was carried out as described in detail in (Binó et al., 2022). Briefly, pOG44 and the corresponding BioID plasmid (CEP89, IFT81, CP110, CEP104, or “no bait control” – BioID plasmid with no insert) were mixed in the ratio of 29:1 (2.9 µg of pOGG44 to 0.1 µg of BioID plasmid per 5 cm culture plate) in OptiMEM (ThermoFisher, Cat. No. 31985070), incubated with Lipofectamine 3000 for 15 minutes, and added drop-wise to the cells. Growth medium was changed to selection medium (G418 (Merck, Cat. No. 4727878001) (400 µg/ml) and Blasticidine (Merck, Cat. No. 203350-25MG) (10 µg/ml)) 24 hours after transfection and was changed every 2-3 days until colonies of resistant cells formed. Expression of the BioID-transgene was induced by doxycycline (Merck, Cat. No. D3072-1ML, see further), and verified by western blotting and/or IF microscopy.

For a rescue experiment, TTBK2 KO cell lines expressing individual BioID-baits were transduced by a lentiviral vector expressing TTBK2 as described in detail in (Binó et al., 2022).

### Zebrafish CRISPR/Cas9 design and sgRNA preparation

Guide-RNA targeting samd4b was designed using the ChopChop web interface (Labun et al., 2019), by. generating candidate gRNA sequences and ranking them based on their predicted efficiency and specificity. The following gRNA sequences (Suppl. Table 3) were chosen for their high probability of introducing a frameshift knockout mutation and minimal potential for off-target cleavage. The selected sequence was integrated by PCR to integrate this sequence into a DNA scaffold using a previously published protocol (Shah et al., 2015). T7 RNA polymerase kit (MEGAscript T7 transcription kit, ThermoFisher Scientific, Cat. No. AM1333) was used to transcribe and purify sgRNAs before resuspending them in nuclease-free H_2_O to a final concentration of 750 ng/ul. Purified sgRNA was then stored in 2 µl single-use aliquots at −80°C.

### Zebrafish breeding, embryo collection and microinjections

Adult wild-type Zebrafish (Tuebingen strain) and the corresponding samd4b heterozygotes (this study) were obtained and processed according to the rules established by the Laboratory Animal Science Committee of Masaryk University (MSMT-34667/2022-5). One day prior to the experiment, males were separated from females. On the day of the experiment, males and females were moved to a shared tank and left to breed for 15-30 minutes. Fertilized eggs laid into the water were collected into a Petri dish with E3 medium (5 mM NaCl, 0.17 mM KCl, 0.33 mM CaCl_2_, 0.33 mM MgCl_2_, in H_2_O). For microinjections, the wild type eggs were gently positioned into individual injection chambers made from 1.5% agarose (ITW Reagents, Cat. No. A8963) dissolved in E3 medium using a Pasteur pipette and a paintbrush. In vitro transcribed sgRNAs were mixed with recombinant SpCas9 protein (Sigma-Aldrich, Cat. No. CAS9PL) and KCl to a final volume of 3 µL (600 ng sgRNA, 2 ug SpCas9, 130 nM KCl, H_2_O to 3 µL) and incubated for 20 min at room temperature to support formation of Cas9 + sgRNA complexes. This mixture was loaded into a glass capillary needle connected to a FemtoJet 4x microinjector (Eppendorf), and 1 nl was injected into individual fertilized eggs, targeting the animal cell-yolk border while they were still in the 1-2 cell stage. The control group was injected with mock solution (SpCas9 and KCl). Injected eggs were gently moved to a fresh Petri dish with E3 medium and left to develop further in a 28.5°C incubator for approximately 60 hours post-fertilization (hpf).

### Genotyping of CRISPR-induced mutations

To generate a stable samd4b mutant line, we outcrossed mosaic F0 founders to wild-type fish and screened the progeny for gamete-transmittable mutations. Briefly, F1 generation fish were anesthetized using tricaine (200 mg/L, Sigma-Aldrich, Cat. No. E10521), a small part of the caudal fin was collected into 20 µL of DNA extraction buffer (25 mM NaOH, 0.2 mM EDTA, in H_2_O) and boiled at 95°C for 10 min to extract genomic DNA. This was repeated for each individual F1 fish, usually in genotyping batches of 30-40 fish where each fish was kept separately after tissue collection. The first genotyping PCR was run using primers closely flanking the CRISPR target site, so that the final PCR product was 90-120 bp long. These short PCR products were then run on a 3% agarose gel (ITW Reagents, Cat. No. A8963) in TBE buffer (100 nM Tris base, 100 nM boric acid, 2 nM EDTA) to detect INDELs as small as 4bp. INDELs that were easy to detect on a gel were selected for further sequencing analysis by a second “NGS” set of primers to amplify a larger genomic region (500-800 bp), which was sequenced using Illumina next-generation sequencing (commercial service) to confirm frame-shift or in-frame mutation. The identified frame-shift samd4b mutant allele (7bpINS) was kept in a heterozygous state following 1-2 additional outcrosses to wild type prior to any experiments. All experiments were carried out by day 5 post-fertilization.

### BioID pulldown

To identify proximity interactions of individual BioID-baits, 2×15-cm plates (high cell density “High ciliated” experiment; 250 cells/mm^2^), or 3×15-cm plates (low cell density “Low ciliated” experiment; 100 cells/mm^2^) of cells expressing each bait were used per 1 sample. 24 hours after seeding, doxycycline (200 ng/ml; Merck, Cat. No. D3072-1ML) was added to the cells. Biotin (50 µM, Santa Cruz Biotechnology, Cat. No. sc-204706) was added the next day for an additional 24 hours. Cells were washed with phosphate-buffered saline (PBS), scraped, and pelleted (200g/5 min). Cell pellets were lysed (all following steps done on ice) in lysis buffer (50 mM Tris-Cl, pH 7.4, 250 mM NaCl, 0.4% SDS, 2% Triton X-100, 1 mM DTT, and 1× Complete proteasome inhibitors (Roche, Cat. No. 4693132001)) for 15 min at 4°C. Following centrifugation (18,000 × g for 15 min at 4°C), cleared extracts were diluted with an equal volume of wash buffer (50 mM Tris-HCl, pH 7.4), incubated (4°C/ON) in an orbital shaker with 20 µl (slurry) Streptavidin sepharose beads (Merck, GE17-5113-01). Bound complexes were pelleted, washed twice with lysis buffer and subsequently twice with wash buffer, transferred to clean tubes, washed once with wash buffer and analysed by MS/MS to define interacting proteins The outlined experiment was carried out at least in two biological replicates. All listed chemicals were purchased from Merck, unless stated otherwise.

### Proteomics

#### On-beads digestion

Bead-bound protein complexes were digested directly on beads by addition of 1 μg/μl trypsin (sequencing grade; Promega) in 50 mM NaHCO_3_ buffer at 37°C with mild agitation for 1 hour. Beads were then vortexed, the supernatant was transferred to clean tubes, and incubated at 37°C for 18 h without agitation. Resulting peptides (∼ 15 μl solution) were extracted into LC-MS vials with 2.5% formic acid (FA) in 50% acetonitrile (ACN) and 100% ACN with the addition of polyethylene glycol (20,000; final concentration 0.001%) and concentrated in a SpeedVac concentrator (ThermoFisher).

#### LC-MS/MS analysis of peptides

LC-MS/MS analyses were performed on an RSLCnano system coupled to an Orbitrap Fusion Lumos Tribrid mass spectrometer (Thermo Fisher Scientific). Tryptic peptides were concentrated and desalted on a trapping column (100 μm × 30 mm, X-Bridge BEH130 C18, 3.5 μm; Waters) and separated on an Acclaim PepMap100 C18 analytical column (75 μm × 500 mm, 3 μm; Thermo Fisher Scientific) using a 100 min nonlinear gradient of 1-56% solvent B (solvent A, 0.1% formic acid in water; solvent B, 0.1% formic acid in 80% acetonitrile) at a flow rate of 300 nL/min. The analytical column was connected to a Digital PicoView 550 ion source equipped with a PicoTip emitter (New Objective), and an active background ion reduction device (ABIRD) was used.

MS data were acquired in data-dependent acquisition mode, selecting the top 10 precursor ions from survey scans (m/z 350-2000). Full MS scans were acquired at a resolution of 60,000 with a target value of 1 × 10^6 ions and a maximum injection time of 200 ms. HCD MS/MS spectra were acquired at a resolution of 15,000 with a target value of 50,000 ions and a maximum injection time of 500 ms. Dynamic exclusion was set to 45 s, and precursor isolation was performed with a 2 m/z window.

Raw data were processed using MaxQuant v1.6.10.43 and the Andromeda search engine. Spectra were first searched against a modified cRAP contaminant database (112 sequences, version 181122) and subsequently against the UniProtKB human proteome database (taxonomy ID 9606; 20,621 entries; downloaded June 17, 2020). Precursor and fragment mass tolerances were set to 10 ppm and 0.05 Da, respectively. Methionine oxidation, asparagine/glutamine deamidation, and protein N-terminal acetylation were specified as variable modifications, and up to two missed cleavages were allowed. Peptide-spectrum matches were filtered using Percolator, and only peptides with q < 0.01, rank 1, and a minimum length of six amino acids were retained. Proteins were reported as protein groups and required at least one unique peptide for identification. Label-free quantification was performed in Proteome Discoverer using the top three peptide approach (Silva et al., 2006).

#### Proximity interactors identification

To search for proteins specifically interacting with individual BioID baits, protein intensity of the BioID “only” control condition was used to filter the primary interaction data. The following criteria were applied for a protein to be considered bait-specific: **A:** The intensity of the protein in pulldowns of BioID-“bait” to BioID “only” was at least 8-fold higher, or the protein was not detected in the Myc-BioID “only” condition; **B:** the protein was detected in two replicates, and once by at least two independent peptides. The proximity interactomes were visualized using Cytoscape (Shannon et al., 2003).

### Microscopy and image analysis

Cells seeded on glass coverslips were fixed in cold MetOH (–20°C/10 min; Fig. 5A-B, Suppl. Fig. 5A-D) or 4%PFA in PBS (RT/10 min; Fig.5F-H, Suppl. Fig. 5H-J, 3× PBS washed (in case of PFA fixation, a permeabilization step with 0.2% Triton X-100 in PBS/10 min/RT was included), blocked (3% bovine serum albumin in PBS with 0.01% NaN_3_), and incubated with primary antibodies (4°C/ON). The antibodies used are listed in Suppl. Table 3. Following 3x PBS washing, the specimens were incubated with secondary antibodies (RT/2 h /dark) and DAPI (1:2000, RT/5 min/dark), washed with PBS 3×, and mounted in ProLong™ Glass Antifade Mountant (ThermoFisher Scientific, Cat. No. P36980).

Microscopy analyses were done using a Zeiss AxioImager Z2 microscope with a Hamamatsu ORCA Flash 4.0 camera, 63× Apo oil immersion objective, and ZEN Blue 2.6 acquisition SW (Zeiss). Image stacks acquired using Zeiss AxioImager.Z2 were projected as maximal intensity images using ImageJ distribution FIJI (Schindelin et al., 2012). All images within the given experiment were acquired with the same laser intensities and exposure times. Where appropriate, contrast and/or brightness of images were adjusted using Photoshop CS5 (Adobe) or FIJI. To evaluate the effect on cilia percentage and cilia length, 5 fields of view containing 60-100 cells were analyzed per condition and experiment by Cilia-Q (Hansen et al., 2021).

Image analysis of signal intensity (Fig. 5A-B, Suppl. Fig. 5A-D) was performed by measuring the centrosome signal intensity of the protein of interest (POI) in FIJI and the corresponding centrosome marker (γTUB) by drawing a circular region of interest (ROI) around the centrosome and measuring mean signal intensities inside the ROI (= centrosome signal), then the background intensity of ROI next to the centrosome (= background signal). The final plotted values are equal to: (centrosome POI signal − background POI) /(centrosome marker signal − background marker signal)

To measure signal intensity in cilia (Fig.5F-H, Suppl. Fig. 5H-J), a segmented line was drawn in FIJI along the primary cilium (defined by signal of ARL13B), and the mean intensity values (in arbitrary units) along the line were recorded. About 30-50 cilia were analyzed per condition and experiment.

Samples for expansion microscopy were prepared as described before (Pejskova et al., 2020) and imaged using the LSM-880 Airy2 microscope (alpha Plan-Apochromat 100x/1.46 Oil DIC M27 Elyra, AiryScan detector) by ZEISS. Raw images (.czi files) were processed with the AiryScan and Orthogonal Projection (maximum intensity) post-processing features in ZenBlack.

Zebrafish embryos (approximately 60 hpf) were gently dechorionated using forceps, anesthetized using tricaine, and placed at 4°C for 15 min. Embryos were then imaged under a brightfield microscope (AxioZOOM, Zeiss); image processing and analysis were done using the FIJI software. Eye size measurements from microscopy images were done by manually labelling eye margins in FIJI, performing an ellipse fit, and measuring the largest ellipsoid axis length.

### RNA isolation and Q-PCR

Zebrafish were anesthetized using tricaine, and caudal fin tissue from 10 individual animals was pooled together into 350 µl of pre-chilled RLT buffer (RNEasy Mini kit, Qiagen, Cat. No.74104) supplemented with 3.5 µl β-mercaptoethanol. The samples were pipetted up and down until the tissue was completely dissolved and then snap-frozen at −80°C for 30 min. After gently thawing the samples on ice, 350 µl of 70% EtOH was added to each sample. The rest of the mRNA isolation was carried out as described in the kit guidelines, including on-column DNase I treatment to degrade residual genomic DNA. Reverse transcription was performed with the Transcriptor First Strand cDNA Synthesis kit (Roche, Cat. No. 04379012001) using a mix of oligo(dT) and random hexamer primers and 600 ng of mRNA as template for each reaction. RNA from NIH 3T3 cells was also isolated using RNeasy Mini Kit (Qiagen, Cat. No. 74106), with 1 ug of RNA used for the cDNA synthesis. Q-PCR reactions were prepared in triplicate using LightCycler 480 SYBR Green I Master Mix (Roche, Cat. No. 04707516001), LightCycler^®^ 480 Instrument II (Roche), and 40-50x diluted cDNA as a template. Relative gene expression was calculated using the 2^-ΔΔCT^ method (Livak and Schmittgen, 2001); mouse Actb (Beta Actin) was used as the housekeeping gene in the NIH 3T3 experiments; hatn10 and dna15ta were used for determining relative expression in zebrafish samples. The primers used are listed in Suppl. Table 3.

### Statistical analysis

Statistical analysis by Student’s t-test or One-way ANOVA (specified in the figure legend) was performed using GraphPad Prism; P < 0.05 (*), P < 0.01 (**), P < 0.001 (***), and P < 0.0001 (****) were considered significant at the indicated level. Results are presented as mean plus SD or SEM, or in the form of a box and whisker plot, where indicated.

## Results

### Selection strategy to identify novel cilia regulators in proteomics datasets

To identify novel regulators of cilia formation and function, we analyzed proximity interactomes for four bait proteins (CEP89, IFT81, CP110, CEP104) and the changes in those in response to the presence or absence of primary cilia in hTERT-RPE1 cells (Fig. 1A). To this end, we compared bait interactomes between wild type (WT) and TTBK2 knockout (KO) cells that fail to initiate cilia formation (Benk Vysloužil et al., 2025; Goetz et al., 2012). The incidence of ciliogenesis was further modulated by a combination of cell density and serum starvation effects, resulting in “High ciliated” (∼ 40 % ciliated WT cells) and “Low ciliated” (∼ 5% ciliated WT cells) conditions (Fig. 1A, Suppl. Fig. 1A). Following the initial validation of the expression and localization of the Myc-BirA (BioID)-tagged baits (Suppl. Fig. 1B), we performed a proximity labeling experiment by streptavidin capture of biotinylated proximity interactors followed by tryptic digest and subsequent LC-MS/MS analysis. WT cell line expressing BioID alone served as a negative control to exclude nonspecific interactors (only hits with signal intensity fold change (fCH) ≥ 8 over the BioID negative control were considered specific).

**Figure 1:**
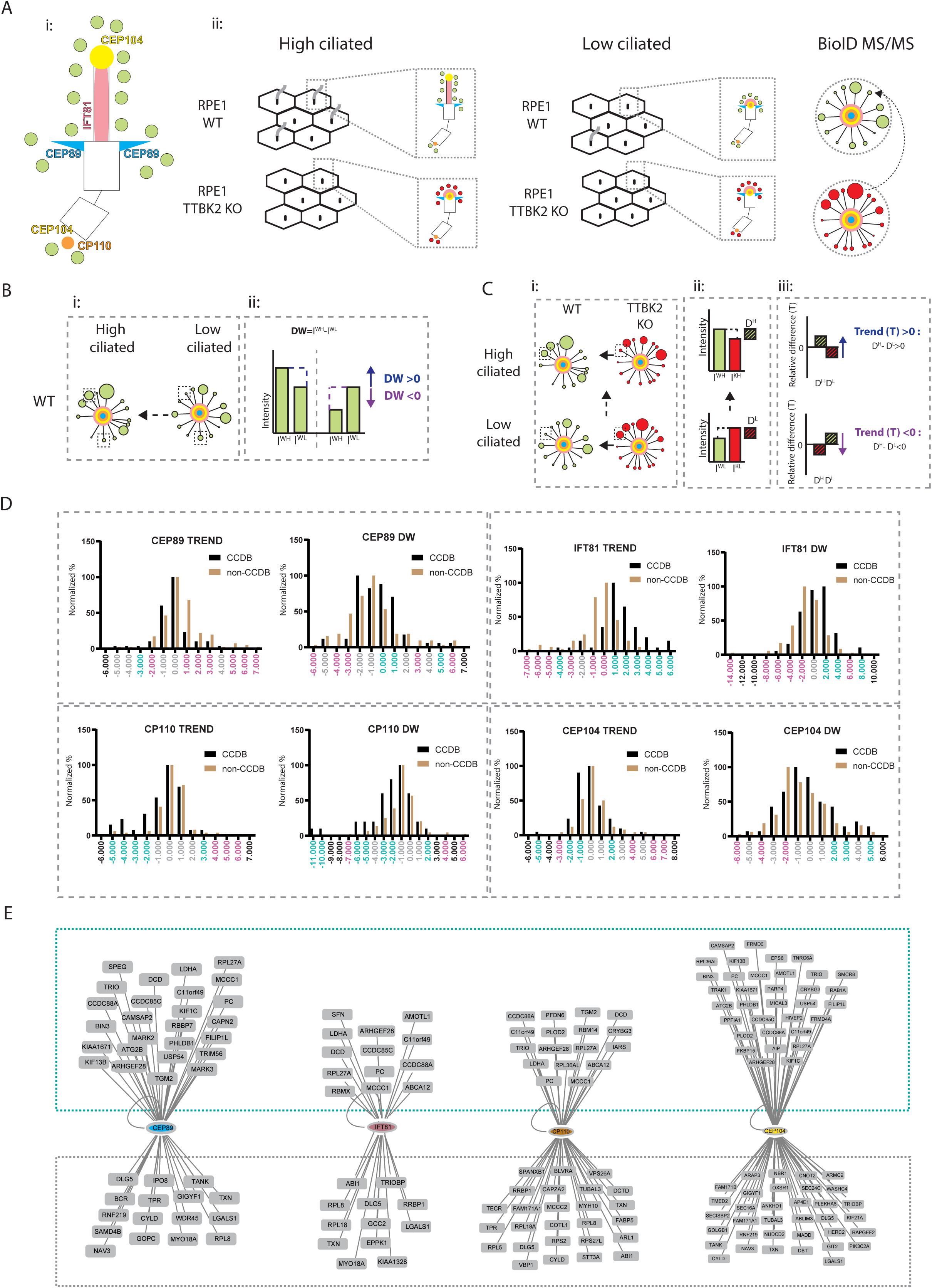
Selection strategy to identify novel cilia regulators in proteomics datasets. **A)** Graphical outline of proximity labeling proteomics experiment. **i:** Sub-ciliary localization of bait proteins (distal appendage component CEP89 is shown in light blue, IFT81 in pink, centriole distal end proteins CP110 and CEP104 in orange, and CEP104 ciliary tip pool in yellow). Green dots represent putative prey proteins. **ii:** Detection of changes in the interactomes of WT and TTBK2 KO cells expressing individual baits under “High ciliated” and “Low ciliated” conditions. Green dots depict prey proteins detected in WT, red dots indicate interactors found in TTBK2 KO for the baits used (center of the interactome). The size of each dot reflects the MS intensity of the corresponding prey protein. **B)** Scheme explaining the use of the Difference (D) parameter, which captures changes between the “High ciliated” and “Low ciliated” datasets for a given condition. **i:** Cartoon depicting potential interactions between bait proteins and prey proteins in WT (green interactome) and TTBK2 KO (red interactome) conditions. The size of each dot reflects the MS intensity of the corresponding prey protein. The arrow indicates that individual prey protein intensities were compared between WT and TTBK2 KO conditions. Dashed line rectangles indicate the protein intensities used as examples in **ii**: I^WH^, I^WL^ represent MS intensity values of example prey proteins (H - “High ciliated”, L - “Low ciliated”), plotted in the graph. A positive DW value (reflecting an increase in protein intensity during ciliogenesis) is indicated by the blue color and the upward arrow, whereas a negative DW value (reflecting a situation where protein intensity drops with ciliogenesis) is indicated by the purple color and the downward arrow. **C)** Scheme explaining the use of the Trend (T) parameter, which uses D values to integrate MS protein intensity in WT cells to that of cells unable to assemble cilia (TTBK2 KO cells), and which also characterizes the behavior of a given protein during ciliogenesis. **i:** Cartoon depicting potential interactions between bait proteins and prey proteins in WT (green interactome) and TTBK2 KO (red interactome) in “High ciliated” (upper panel) and “Low ciliated” (lower panel) conditions. The size of each node reflects the MS intensity of the corresponding prey protein. The arrows indicate the comparisons of individual prey protein intensities between the indicated conditions. Dashed rectangles mark the protein intensities used as examples in **ii:** Graphs illustrate how D^H^ and D^L^ values are calculated. Here, I^WH^, I^KH^, I^WL^, and I^KL^ represent MS intensity values of example prey protein (H-“High ciliated”, L - “Low ciliated”, W - WT, and K - TTBK2 KO). **iii**: Trend (T) represents the difference between D^H^ and D^L^. Positive T value (∼typically reflecting an increase in protein abundance within a particular interactome during induction of ciliogenesis and in WT relative to TTBK2 KO cells) is indicated by blue color and the upward arrow, whereas negative T value (∼ typically reflecting a decrease under these conditions) is shown in purple color and downward arrow. **D)** Stratification of identified prey proteins according to their T/DW values. Normalized values (normalized percentage) for corresponding T and DW values are plotted on the Y axis. The X-axis shows T or DW values. The dashed boxes group T- and DW-based graphs for each bait. Prey proteins annotated as members of CCDB are indicated by black bars, whereas non-CCDB proteins are indicated by brown bars. “Enriched” groups (containing at least 30% more CCDB than non-CCDB proteins) are indicated in turquoise; “depleted” groups (containing at least 30% fewer CCDB proteins than non-CCDB proteins) are indicated in magenta. “Neutral” groups are shown in grey. **E)** Cytoscape-based interactomes of used bait proteins (depicted as the central node of each interactome) comprising non-CCDB proteins sorted as either “Enriched” (indicated by turquoise dashed line box) or “Neutral” (indicated by grey dashed line box) by analyses outlined in D.

Increased signal intensity of prey protein reflects the increased proximity interactions with the used bait protein (Firat-Karalar and Stearns, 2015; Iazzi et al., 2024; Kim and Roux, 2016). Indeed, we observed intensity changes for many prey proteins we detected, indicating significant remodeling of protein complexes in relation to cilia formation, as expected. However, comparing the intensity changes of individual prey proteins across different baits (note the selected examples such as MPHOSPH9, CEP152, RPGRIP1L, etc. in red (Suppl. Fig. 1C)) did not reveal any obvious trends in interaction changes of the examined proteins across the baits.

Building on the fact that protein abundance is proportional to the abundance of its peptides and, in turn, to the sum of the signal intensities of those peptides measured by MS analysis (Vogel and Marcotte, 2009), we established peptide intensity-derived parameters to assess changes across different conditions and baits. The first parameter, termed “Difference” **D**, captures changes between the “High ciliated” and “Low ciliated” datasets for a given condition. Specifically, we used **DW** to determine the effect of ciliogenesis (changes between “High ciliated” and “Low ciliated”) in the WT condition (Fig. 1Bi-ii). The second parameter, termed **“Trend” T**, uses DW values to integrate MS intensity of a given protein in WT cells to that of cells unable to form cilia (TTBK2 KO cells, Fig. 1Ci-ii). It also characterizes the behaviour of a given protein during ciliogenesis by comparing its relative abundance between “High ciliated” and “Low ciliated” datasets (Fig. 1Ciii). A positive T value reflects an increase in protein abundance within a particular interactome during ciliogenesis and in WT relative to TTBK2 KO cells, whereas a negative T value indicates a decrease in protein abundance under these conditions (Fig. 1Ciii).

We found 375 proteins common in both “High ciliated” and “Low ciliated” experiments, with 193 proteins detected for more than 1 bait (Suppl. Table 1). Next, we defined a “CCDB (Centrosome and Cilium Database)” category in our datasets as proteins already annotated as centrosome/cilia components (Gupta et al., 2015). For CEP89, we identified 70 CCDB proteins compared to 121 non-CCDB proteins; for IFT81, 61 CCDB and 102 non-CCDB; for CP110, 42 CCDB and 122 non-CCDB; and for CEP104, 67 CCDB and 129 non-CCDB proteins (Suppl. Table 1). Next, we stratified these proteins according to their T and DW values into subgroups (Fig. 1D). We postulated that putative novel cilia/centrosome regulators among the non-CCDB proteins would be more likely to cluster with CCDB proteins. Based on the abundance of CCDB proteins (reflected by the Normalized % values on the Y-axis in Fig. 1D), we classified each subgroup as “Enriched,” “Depleted,” or “Neutral”. A subgroup was considered “Enriched” if it contained at least 30% more CCDB than non-CCDB proteins, and “Depleted” if it contained at least 30% more non-CCDB than CCDB proteins. Subgroups falling between these thresholds were considered “Neutral.” Intriguingly, we observed distinct patterns in the T/DW distribution of prey proteins for the IFT81 and CP110 baits. For IFT81, the prey proteins with positive T/DW values were typically found “Enriched”, while groups with negative T/DW belonged to the “Depleted” category (Fig.1D, compare black to brown bars). This is in line with the current understanding of the assembly and positive function of IFT complexes in cilia formation (Lacey and Pigino, 2025). In the case of CP110, the “Enriched” prey proteins often had negative values for both DW and T, consistent with the loss of CP110 and its interactions from the basal body at the onset of ciliogenesis (Gupta et al., 2015; Spektor et al., 2007). No obvious grouping was observed for CEP89 and CEP104 data.

Next, we determined interactomes comprising “Enriched/Neutral” non-CCDB proteins for each bait (Fig. 1E). For CEP89, 29 proteins were classified as “Enriched” and 15 as “Neutral”. The IFT81 interactome contained 16 “Enriched” and 13 “Neutral” proteins, whereas the CP110 interactome comprised 25 “Enriched” and 28 “Neutral” proteins. The CEP104 interactome contained 39 “Enriched” and 35 “Neutral” proteins. Intriguingly, several non-CCDB proteins labelled “Enriched” or “Neutral” have been recently identified as novel cilia/centrosome components (e.g., ARMC9 (Van De Weghe et al. 2017), C11ORF49 (Gheiratmand et al., 2019), KIAA1328 (Drew et al., 2017), KIF13B (Rezi et al., 2025; Schou et al., 2017), CCDC88A/Girdin (Nechipurenko et al., 2016) or CYLD (Douanne et al., 2019). We concluded that this reflects the significant progress made since the CCDB proteins were originally reported (Gupta et al., 2015). Importantly, it has also implied that the data stratification pipeline has the potential to point out putative regulators of centrosome/ciliogenesis among non-CCDB proteins.

### Rescue of TTBK2 function restores the proximity interaction landscape

To further evaluate the functionality of the selection strategy, we analyzed interactome changes in TTBK2 KO RPE-1 cells with restored TTBK2 activity. First, following lentiviral transduction of Flag-TTBK2 into TTBK2 KO BioID-bait (-CEP89-, -IFT74, -CP110, -CEP104) cell lines, we confirmed that expression of the kinase restored primary cilia formation, as expected (Suppl. Fig. 2A). In turn, we carried out the proximity labeling experiment using “High ciliated” and “Low ciliated” protocols. To assess the extent of rescue, we determined additional “Difference” parameters: DR and DK to compare changes across individual datasets (Fig. 2A). Specifically, we focused on proteins common between the DW/T datasets (Fig.1) and Rescue datasets that exhibited either positive (T> 0.7) or negative (T<−0.7) Trend values (Suppl. Table 2). Intriguingly, we observed a shift in D values for many such proteins in the rescue experiment (DR values were often more similar to DW than to DK, particularly in the CEP89 and IFT81 interactomes (Fig. 2B, Suppl. Table 2). This suggests that TTBK2 rescue cells resembled WT cells more than parental TTBK2 KO, owing to the partial rescue effect of TTBK2 reintroduction on the proximity interactions of the examined proteins.

**Figure 2:**
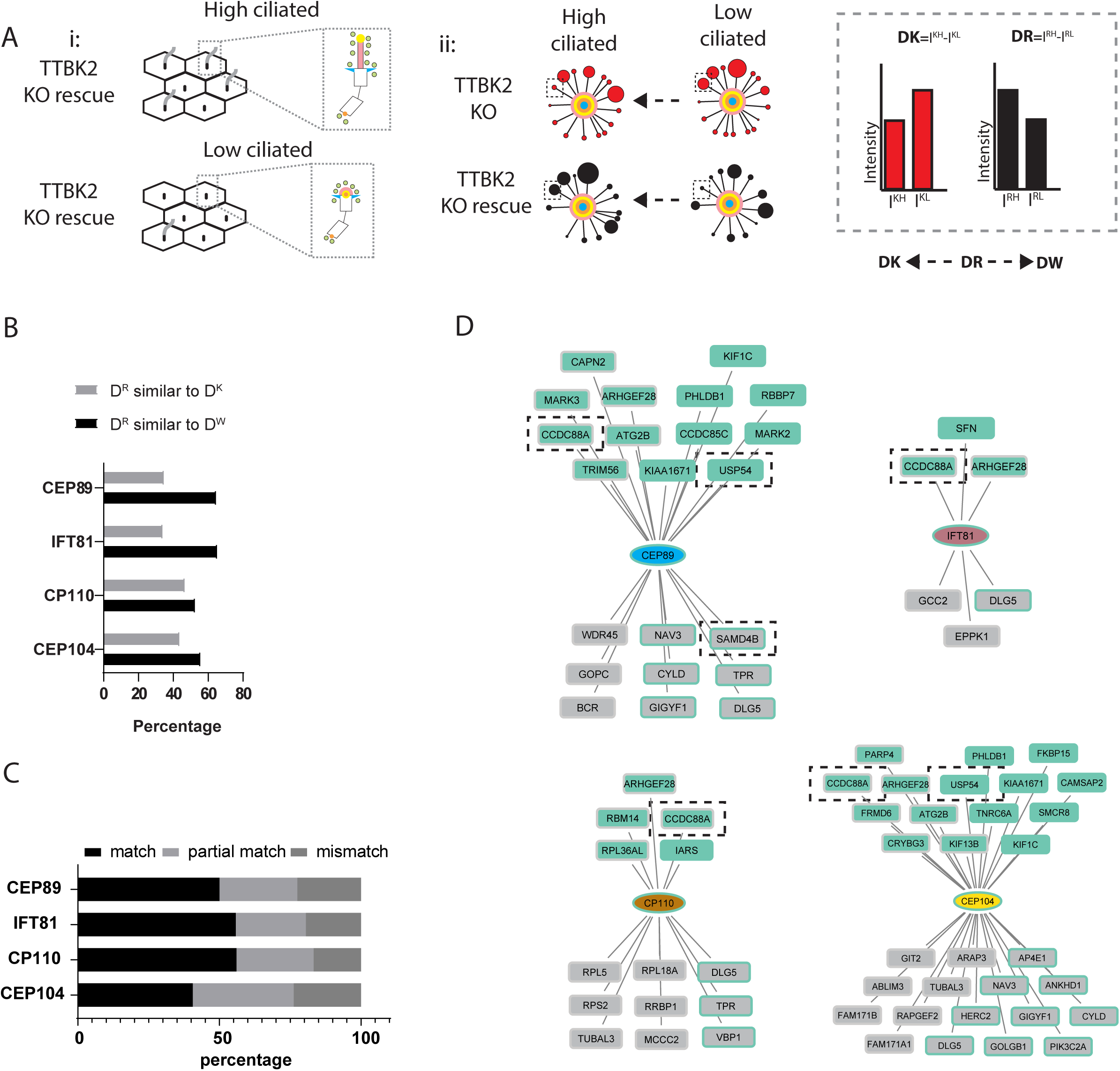
Rescue of TTBK2 function restores the proximity interaction landscape. **A)** i: Graphical outline of proximity labeling proteomics rescue experiment using TTBK2 KO re-expressing exogenous TTBK2 (TTBK2 KO rescue). ii: Detection of changes in the interactomes of TTBK2 KO (Fig.1) and TTBK2 KO rescue cells expressing individual baits under “High ciliated” and “Low ciliated” conditions to determine “Difference” parameter DK and DR. The interactomes depict proximity interactions between bait proteins and prey proteins in TTBK2 KO (red interactome) and TTBK2 KO rescue (black interactome). The size of each dot reflects the MS intensity of the corresponding prey protein. The extent of rescue was assessed by comparing DR to DW (determined in Fig.1) and DK. **B)** Analysis of changes in protein proximity interactions detected in TTBK2 KO rescue cells (TTBK2 KO cells re-expressing exogenous TTBK2) based on DR, DW, and DK comparison. The graph shows the percentage of proteins in each interactome whose DR values were similar to DW (black bars) or to DK (grey bars). **C)** Overlap between the “Enriched”, “Neutral”, and “Depleted” categories based on the T/DW and TTBK2 KO rescue datasets. Proteins independently assigned to identical categories are labelled as “match” (black), whereas those assigned to opposite categories (“Enriched” versus “Depleted”) are marked as “mismatch” (dark grey). The “partial match” (light grey) group includes proteins classified as “Neutral” in the T/DW analyses but as “Enriched” or “Depleted” in the TTBK2 KO rescue dataset, or vice versa. **D)** Cytoscape-based visualization of non-CCDB proteins classified as “Enriched” or “Neutral” for each bait (CEP89 - light blue, IFT81 - pink, CP110 - orange, or CEP104 - yellow). Proteins in the “Enriched” category are shown in turquoise, and those in the “Neutral” category are shown in grey. The sorting based on the T/DW analyses is indicated by the color of the node interior, whereas the TTBK2 KO rescue dataset-based classification is represented by the color of the node border. Dashed rectangles indicate proteins selected for follow-up analysis.

Next, we used DR as a sorting parameter to obtain the “Enriched”, “Neutral”, and “Depleted” categories for the Rescue dataset (Suppl. Fig. 2B), similarly to what we did before using the DW/T datasets (Fig.1). We then compared the overlap between DR-based sorting and the sorting carried out based on T/DW parameters. A fraction of proteins (15-25% of proteins, depending on the bait) that were sorted into the opposite categories formed a “mismatching” group. A larger group (about 25-30% of proteins for each bait) comprised proteins labeled as “Neutral” by one approach, while either “Enriched” or “Depleted” by the other (forming a “partially matching” group). Importantly, the majority of proteins (∼50% of prey proteins per bait) were sorted into the same category (in turn termed “matching”) by both T/DW-based and DR-based sorting (Fig. 2C), further supporting our conclusion that re-expression of TTBK2 in TTBK2 KO cells partially rescued examined protein interactions.

Finally, we used the TTBK2 KO rescue dataset to further narrow down the search for putative novel non-CCDB cilia regulators. To this end, we focused on interactions detected across T/DW and TTBK2 KO rescue datasets and at the same time consistently sorted into either “Enriched” or “Neutral” categories (Fig. 2D, Suppl. Fig. 2C). By combining the results with a literature search, we selected three candidates to focus on: CCDC88A, USP54 and SAMD4B (Fig. 2D). In addition, we included two poorly characterized CCDB proteins into our analysis, CCT3 and BLTP1 (also known as KIAA1109) (Suppl. Table 1).

### SAMD4B and BLTP1 are novel regulators of primary cilia and the Shh pathway

To assess the potential role of the selected candidate proteins in ciliogenesis, we first used siRNA to knock down their expression in hTERT-RPE1 cells. TTBK2 depletion was used as a positive control. After confirming efficient depletion of the individual mRNAs (Suppl. Fig. 3A), we examined the formation of ARL13b-positive primary cilia under each condition. Depletion of CCT3, USP54, SAMD4B, or BLTP1 did not alter the proportion of ciliated cells, in contrast to TTBK2 depletion (Fig.3A and 3B). In the case of CCDC88A depletion, we observed a modest increase in the proportion of ciliated cells. Interestingly, we noted effects on cilia length: primary cilia were shorter following CCT3 and SAMD4B depletion, respectively, whereas knockdown of BLTP1 increased primary cilia length (Fig. 3A, Fig. 3C).

**Figure 3:**
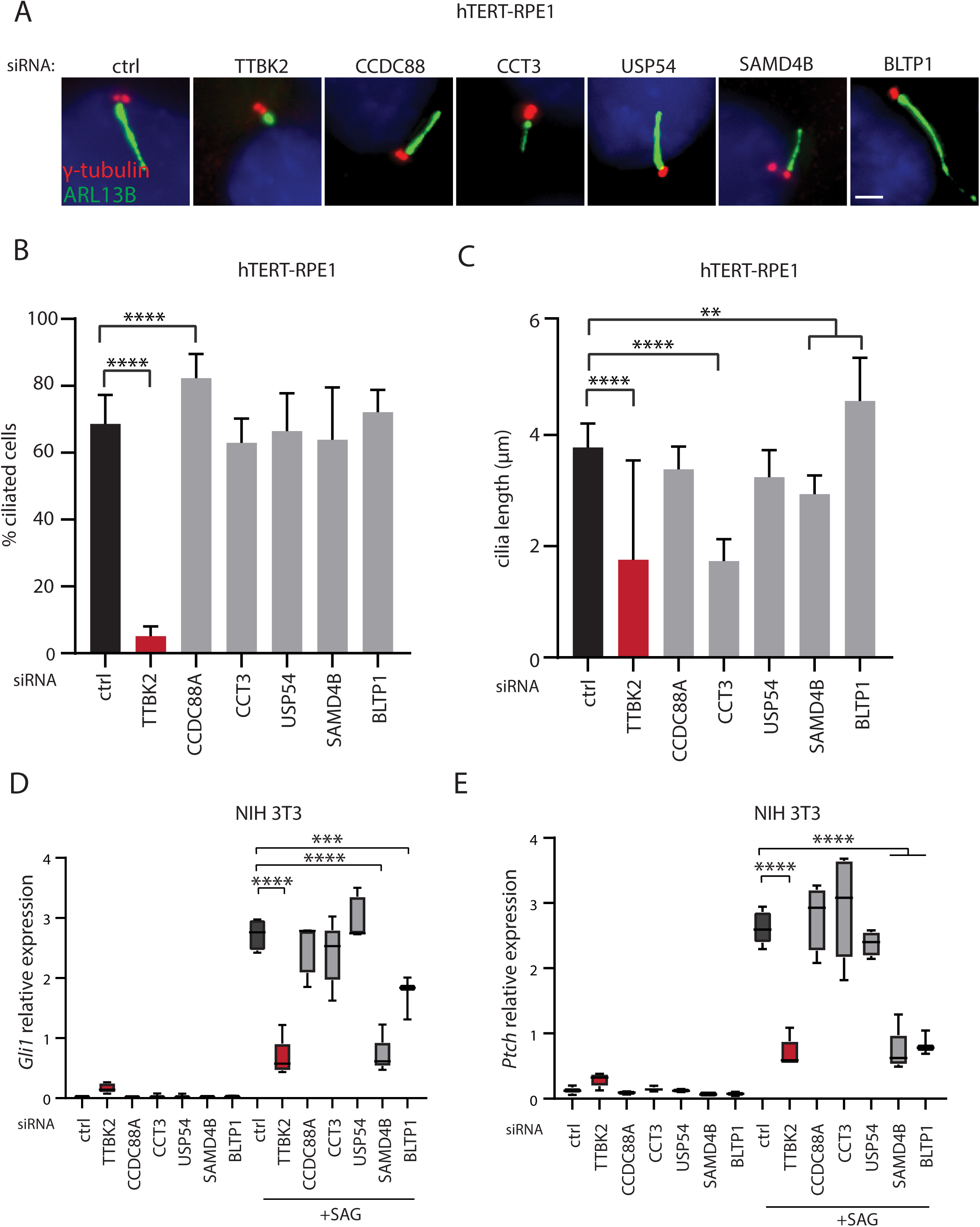
SAMD4B and BLTP1 are novel regulators of primary cilia and the Shh pathway. **A-C)** hTERT-RPE1 cells, transfected with the indicated siRNA, were fixed and stained for ARL13B (green) to visualize cilia and γ-tubulin (red) to detect centrosomes. DAPI (blue) was used to counterstain nuclei. Representative images are shown in **A)**, scale bar 1.5 µm. **B)** Quantification of the percentage of ciliated cells in the indicated conditions. The graph represents a summary of three independent experiments, with at least 200 cells analyzed in each experiment, with a mean and SD shown. **** p<0.0001 by one-way ANOVA. **C)** Quantification of the primary cilium length in the indicated conditions. The graph represents a summary of three independent experiments, with at least 200 cells analyzed in each experiment, with a mean and SD shown. **p<0.01, **** p<0.0001 by one-way ANOVA. Q-PCR analyses of mRNA expression of Shh target genes *Gli1* **D)** and *Patched* **E)** in NIH 3T3 cells transfected with the indicated siRNA and treated with vehicle or 200 nM SAG. Each box-and-whisker graph represents a summary of at least three independent experiments. *** p<0.001, **** p<0.0001 by one-way ANOVA.

To examine a possible role of tested candidates in the cilia-mediated Shh signaling pathway, we turned to NIH 3T3 cells. First, we verified siRNA knockdown efficiency for our candidate genes and TTBK2 (Suppl. Fig. 3B). While depletion of none of our tested candidates affected the percentage of NIH3T3 cells bearing a primary cilium (Suppl. Fig. 3C), depletion of either SAMD4B or BLTP1 abrogated the induction of mRNA expression of Shh pathway target genes *Gli1* and *Ptch* following treatment with 200 nM SAG (Fig. 3D-E). In sum, our data suggest that while SAMD4B, CCT3, and BLTP1 modulate primary cilia length in hTERT-RPE1 cells, only SAMD4B and BLTP1 are required for the induction of the Shh pathway target genes in NIH 3T3 cells, indicating cell-type-specific effects.

### Loss of Samd4b leads to microphthalmia and inner ear defects in vivo

To examine the role of our candidates in vivo, we turned to the zebrafish embryo as a model system. First, we injected gRNA targeting *samd4b*, *bltp1*, or *smo* (used as a positive control) together with recombinant Cas9 into 1-2 cell stage embryos. The ability of the used gRNA to generate frame shift mutations was confirmed by DNA sequencing (Suppl. Fig. 4A). Embryos injected with *smo* gRNA showed notably smaller eyes at about 60 hpf (Fig. 4A-B), in line with the established role of Shh signaling pathway in vertebrate eye development (Chassaing et al., 2016; Ginocchio et al., 2008; Schimmenti et al., 2003; Stenkamp et al., 2002). Intriguingly, we observed a comparable reduction in the eye size at about 60 hpf in embryos injected with *samd4b* gRNA (Fig.4A-B), but not in those injected with *bltp1* gRNA (Suppl. Fig. 4B-C).

**Figure 4:**
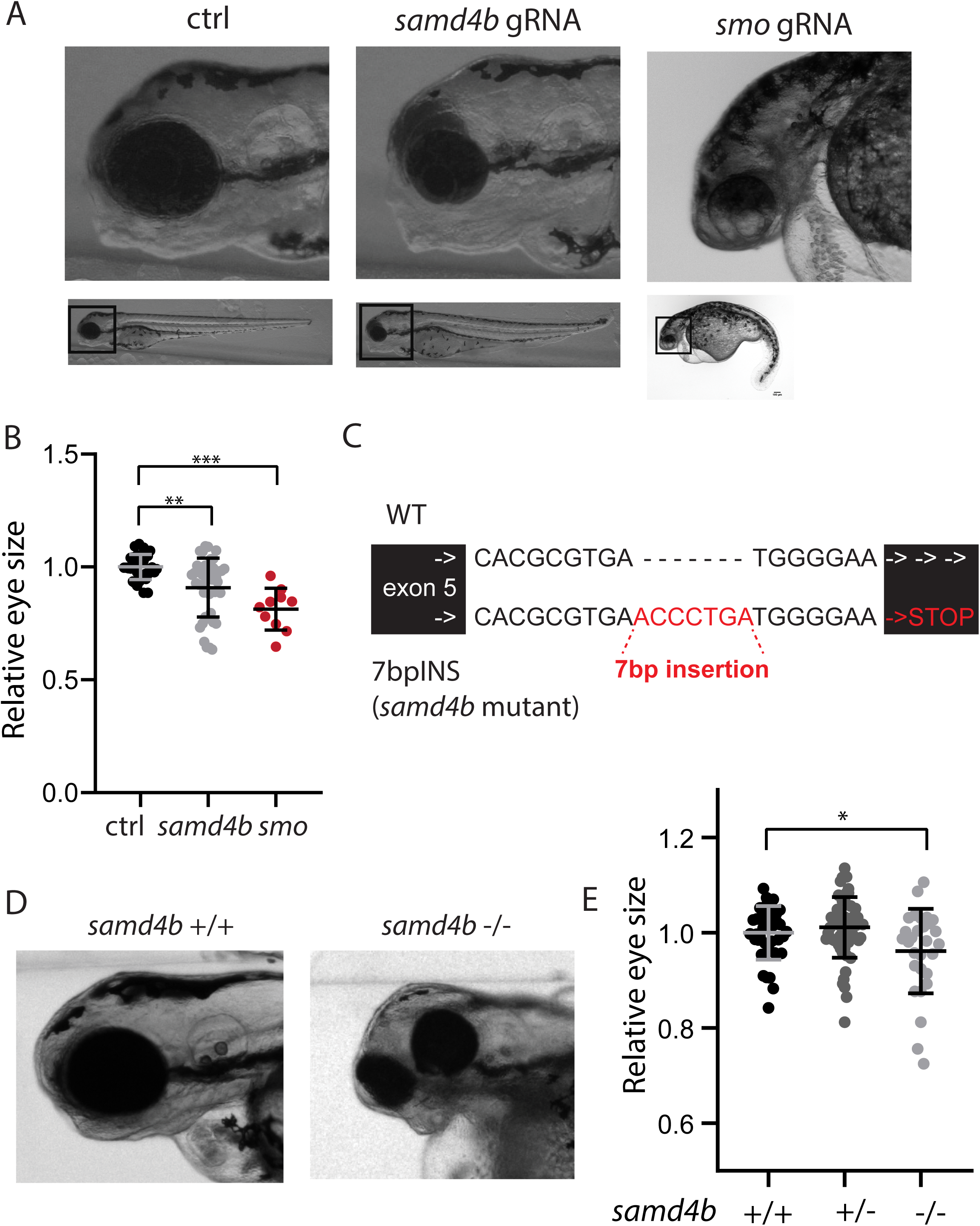
Loss of Samd4b leads to microphthalmia in vivo. **A-B)** Zebrafish embryos were injected with the indicated gRNA or mock and analyzed at about 60 hpf stage. Microphotographs illustrating reduced eye size after *samd4b* gRNA or *smo* gRNA injection are shown in **A).** The graph in **B)** is a summary from four independent experiments and shows relative eye size (related to the control), with each datapoint representing the analyzed fish in the indicated condition, with a mean and SD shown. ** p<0.01, *** p<0.001 by one-way ANOVA. **C)** Schematic illustration of the *samd4b* mutant allele (7bpINS) with insertion of 7 nucleotides into exon 5 of the *samd4b* locus. **D-E)** Zebrafish larvae of the indicated genotypes were analyzed at about 60 hpf. **D)** Microphotographs illustrating reduced eye size in *samd4b* −/−. **E)** The graph is a summary from three independent experiments and shows relative eye size (related to the control), with each datapoint representing the analyzed fish in the indicated condition, with a mean and SD shown. * p<0.05 by one-way ANOVA.

Analysis of crispants from the F0 generation allows very rapid screening for phenotypes, with trade-offs such as chimerism and possible off-targets (Wang et al., 2022). To confirm and extend our initial observations, we generated *samd4b* mutant fish with a 7 bp frame shift insertion in the exon 5 (we termed the mutant allele 7bpINS) (Fig. 4C, Suppl. Fig.4D). As frame shift mutations, and, in turn, the premature stop codon, can lead to degradation of the mutant mRNA by the nonsense-mediated decay (NMD) mechanism (Nickless et al., 2017), we tested whether this applies to *samd4b* 7bpINS mRNA. Indeed, our Q-PCR analyses of *samd4b* mRNA levels showed a marked reduction of the mRNA in *samd4b* +/7bpINS fish (F2 generation) over *samd4b* +/+ fish, suggesting that mRNA containing 7bpINS is indeed efficiently degraded by NMD (Suppl. Fig. 4E) and hence, the 7bpINS allele is likely a complete loss-of-function mutation (we use “-” to indicate this allele from now on). Next, we crossed *samb4b* +/-fish till F3 generation. The obtained offspring in F3 comprised approximately 30 % +/+, 50 % +/-, and 20 % −/− *samd4b* fish, in line with expected Mendelian ratios (Suppl. Fig. 4F). Importantly, we observed that the eyes of *samd4b* −/− fish were moderately smaller than those of *samd4b* +/+ or *samd4b* +/- fish (Fig. 4D-E), in line with our findings from *samd4B* F0 crispants.

Our analysis of *samd4b* mutant fish did not reveal a signature defect in body curvature, typical for mutations in *smo* and several key cilia regulators in zebrafish and leading to ventrally curved fish (Ingham, 2022; Zhang et al., 2018). This suggested that the absence of fully functional Samd4b is either compensated (Rouf et al., 2023) and/or that the role of Samd4b in zebrafish is cell type-specific. To test the latter possibility, we searched for additional phenotypes with established causal links to the Shh pathway and cilia dysregulation. One such phenotype relates to the inner ear, where the residing cilia tether biomineralized aggregates termed the otoliths. In turn, a defect in otolith formation serves as evidence of inner ear cilia dysfunction and is found with variable penetrance in almost all zebrafish ciliary mutants (Whitfield, 2019). Interestingly, *samd4b* −/− fish displayed a trend of higher incidence of otolith abnormalities when compared to *samb4b* +/+ and *samd4b* +/- fish (Suppl. Fig. 4G-H). Importantly, we observed a similar trend when re-examining the F0 *samd4b* crispants (Suppl. Fig. 4I). In sum, our data suggest that Samd4b is involved in the development of the eye and inner ear, hence mimicking some of the Shh- and cilia-related defects in the fish. Further work will be necessary to resolve the underlying mechanism and the anticipated connection between Samd4b and the Shh pathway in zebrafish.

### SAMD4B localizes to centrioles and is required for ciliary localization of SMO and ARL13B

SAMD4B, together with its paralog SAMD4A, is related to a Drosophila protein Smaug, implicated in RNA binding and associated with cytoplasmic RNA granules (Dahanukar et al., 1999; Luo et al., 2010; Youn et al., 2018). Intriguingly, recent proteomics revealed numerous interactions between RNA granule proteins and components of the centrosome, implying an unexpected connection between these two organelles (Youn et al., 2018).

To investigate SAMD4B mechanisms related to cilia and Shh pathway regulation, we examined its ability to localize to centrosomes or cilia in hTERT-RPE1 and normal human dermal fibroblasts (nHDF). While we did not reliably detect the SAMD4B signal inside cilia, we observed it at the centrosome and/or at the cilium base, often associated with either one or both centrioles (Fig. 5A and 5C). We confirmed the specificity of the staining by depleting SAMD4B protein levels with siRNA (Fig. 5A-B) and by using another SAMD4B antibody (Suppl. Fig. 5A-B). To confirm and extend the data on centrosomal localization of SAMD4B, we transfected hTERT-RPE1 cells with a SAMD4B expression vector and observed an elevation of SAMD4B levels associated with the centrosome (Suppl.Fig. 5C-D).

**Figure 5:**
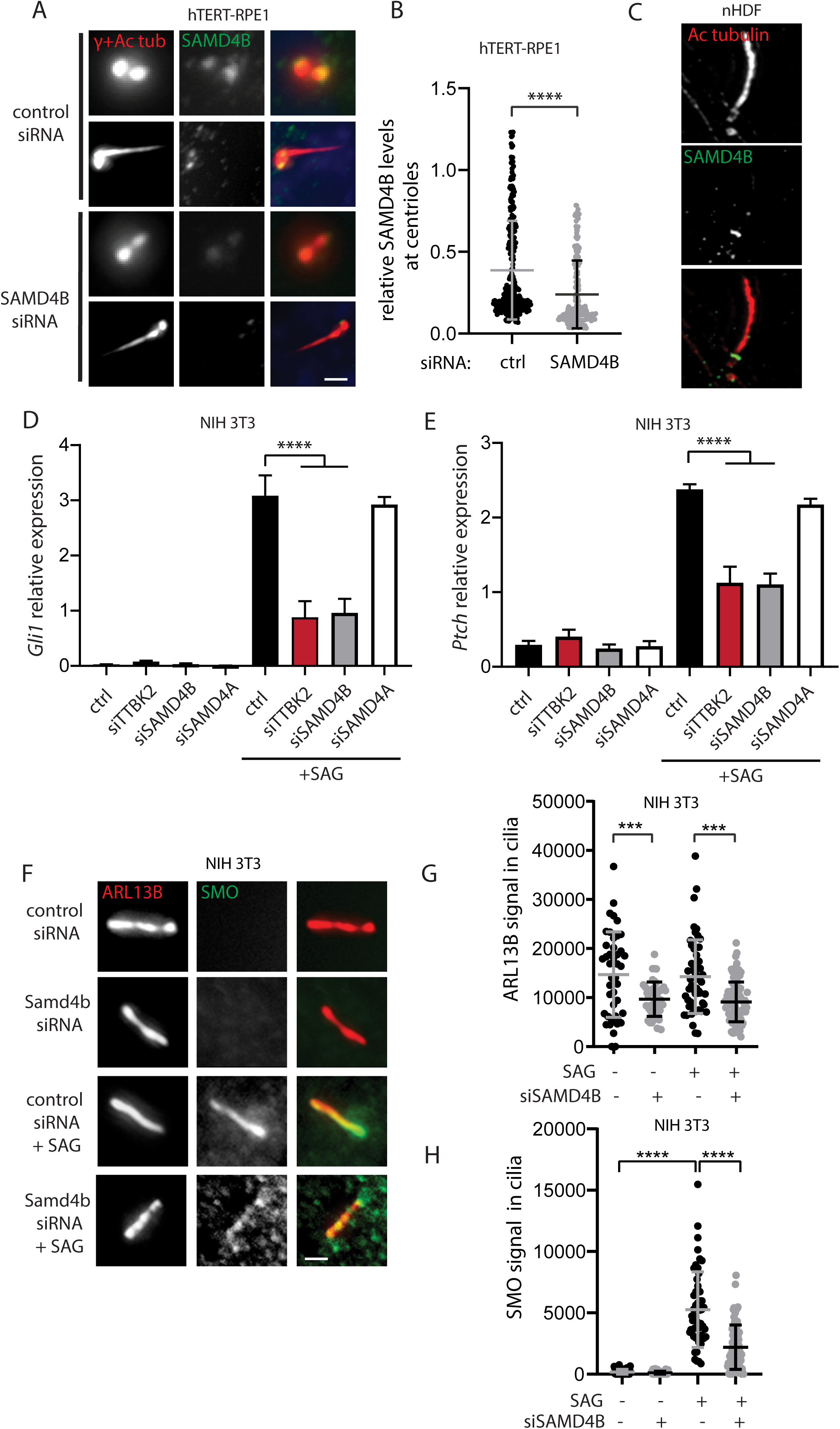
SAMD4B localizes to centrioles and is required for SMO and ARL13B ciliary localization. **A-B)** hTERT-RPE1 cells, transfected with the indicated siRNA, were fixed and stained for SAMD4B (green; Sigma antibody) and acetylated tubulin and γ-tubulin (red) to visualize cilia and centrioles. Representative images are shown in **A),** scale bars 1.5 µm (cilium) and 0.5 µm. Quantification of the relative SAMD4B signal intensity at centrioles (signal of SAMD4B normalized to signal of acetylated tubulin and γ-tubulin) is shown in **B).** The graph represents a summary of three independent experiments, with individual datapoints, mean, and SD shown. **** p<0.0001 by two-tailed t-test. **C)** Representative images of SAMD4B (green; Sigma antibody) and acetylated tubulin (red) staining in nHDF cells, analyzed using expansion microscopy. Q-PCR analyses of mRNA expression of Shh target genes *Gli1* **D)** and *Patched* **E)** in NIH 3T3 transfected with the indicated siRNA and treated with vehicle or 200 nM SAG. Each graph represents a summary of three independent experiments, with mean and SEM shown. **** p<0.0001 by one-way ANOVA. **F-H)** NIH 3T3 cells, transfected with the indicated siRNA and treated with vehicle or 200 nM SAG, were fixed and stained for SMO (green) and ARL13B (red). Representative images are shown in **F)**. Scale bar 1.5 µm. Quantification of the ARL13B signal intensity in cilia is shown in **G)**. Quantification of the SMO signal intensity in cilia is shown in **H)**. Each graph represents a summary of two independent experiments, with individual datapoints, mean, and SD shown. **** p<0.0001, *** p<0.001 by one-way ANOVA.

Next, to corroborate our findings on the requirement of SAMD4B for SAG-induced Shh target genes expression, we depleted SAMD4B using three individual siRNAs (note that the original results in Fig. 3D-E were obtained following SAMD4B depletion using a mix of siRNA#1 and #2). Importantly, each individual siRNA (siRNA#1, siRNA#2, and siRNA#3) depleted *Samd4b* mRNA levels and, in turn, reduced *Gli1* and *Ptch* mRNA expression in SAG-treated NIH 3T3 cells (Supp. Fig. 5E). In addition, we asked whether the protein paralogous to SAMD4B, SAMD4A, is also involved in the regulation of Shh pathway target genes expression. Interestingly, we found that depletion of SAMD4B, but not SAMD4A, impaired SAG-induced mRNA expression of *Gli1* and *Ptch* (Fig. 5D-E, Suppl. Fig. 5F), indicating that the role of SAMD4B in Shh pathway regulation is specific and non-redundant with SAMD4A.

To pinpoint the mechanism responsible for the reduced response of SAMD4B-depleted cells to Shh pathway activation, we examined ciliary levels of components crucial for cilia signaling capabilities, such as a Shh pathway receptor SMO (Corbit et al., 2005; Huangfu and Anderson, 2005) and a membrane-associated small GTPase of the ARF/ARL family – ARL13B (Caspary et al., 2007; Cevik et al., 2010; Larkins et al., 2011). Importantly, depletion of SAMD4B in NIH 3T3 cells reduced ARL13B ciliary levels in both steady state conditions and following Shh pathway activation by SAG (Fig. 5F-G), and impaired SMO ciliary accumulation in the SAG-treated cells (Fig. 5F and 5H). In addition, we detected a similar reduction in ARL13B and SMO ciliary levels also in nHDF cells depleted of SAMD4B (Suppl. Fig. 5G-J). In sum, these results suggest that SAMD4B is required for proper Shh pathway activation through regulation of ciliary levels of ARL13B and SMO.

Conclusively, our data represent the first report describing the role of SAMD4B in the Shh signaling pathway in vertebrate cilia.

## Discussion

Data presented herein identify SAMD4B and BLTP1 as novel centrosome/cilium-associated factors regulating the Shh signaling pathway, with *samd4b* mutant showing defects reminiscent of Shh pathway deregulation also in zebrafish.

Our candidate proteins were revealed in a proteomics screen, comparing proximity interactome changes in ciliated (TTBK2 WT) and non-ciliated (TTBK2 KO) cells, under ciliogenesis-permissive and restrictive cell densities. As expected, many prey proteins exhibited substantial intensity changes, indicating extensive remodeling of protein interaction networks during ciliogenesis, in line with previous work (Gupta et al., 2015). No consistent trends emerged when comparing intensity changes of individual prey proteins across the different baits. This is likely owing to the complexity of interactions occurring during ciliogenesis (Gupta et al., 2015; May et al., 2021) and the specific, non-overlapping roles served by the proteins used as baits.

SAMD4B, together with SAMD4A, are the two vertebrate orthologs of the Drosophila protein Smaug, an RNA-binding protein (in a SAM domain-dependent manner) and repressor of translation (Baez and Boccaccio, 2005; Dahanukar et al., 1999; Luo et al., 2010; Smibert et al., 1999). Given that, the association of SAMD4B with the centrosome and ciliary signaling may seem somewhat surprising. Importantly, however, SAMD4B was detected earlier as a proximity interactor of distal appendage component CEP89 or centrosomal protein CEP152 in HEK 293T cells (Gupta et al., 2015). Moreover, our results are in agreement with recent notions of numerous interactions between RNA-binding proteins and components of centrosomes or cilia (Pachinger et al., 2025; Youn et al., 2018). Given that we found using IF microscopy that SAMD4B is associated with either one or both centrioles, we speculate that this heterogeneity may reflect the dynamic nature of SAMD4B association with centrosomes, the state of centrosome maturation, or the existence of distinct protein pools of SAMD4B residing in this compartment.

Furthermore, our results indicate that proper Shh signaling requires SAMD4B, but not SAMD4A, at least in NIH 3T3 cells. This implies that although both proteins show high sequence similarity, including the SAM domain, SAMD4B seems to display a non-redundant function. In fact, both proteins are considered to have distinct expression patterns, including markedly different expression in tumors, further indicative of possibly divergent functions (Wang and Zhang, 2023).

Our data further suggest that SAMD4B regulates the Shh pathway at the level of SMO and ARL13B ciliary recruitment. Indeed, defective ciliary localization of either of these proteins leads to abrogated Shh signaling in various model systems (Bay et al., 2018; Corbit et al., 2005; Shi et al., 2023). In Drosophila, Smaug directly interacts with SMO to promote Shh signaling in the developing wing (Bruzzone et al., 2020). Thus, direct interaction of SAMD4B and SMO could, at least in principle, explain the effects of SAMD4B depletion on SAG treatment-induced ciliary enrichment of SMO. However, we consider this possibility not very likely, given that Shh signaling in Drosophila is cilia-independent (Ingham, 2022) and that the SMO binding interface of Smaug does seem to be conserved in the vertebrate SAMD4B (Kubíková et al., 2023). Future work should address the exact role of SAMD4B in the regulation of SMO and ARL13B ciliary levels.

Samd4b deficiency in zebrafish leads to reduced eye size and, to some extent, to a higher incidence of otolith abnormalities, both phenotypes commonly found in cilia/Shh pathway mutants in zebrafish (Ingham, 2022; Stenkamp et al., 2002; Whitfield, 2019). This observation supports a model in which reduced Shh signaling causes eye size defects owing to reduced cell proliferation and/or survival (Stenkamp et al., 2002). As we further found no indication of left-right symmetry defects or ventrally curved fish, also typical for many cilia mutants (Ingham, 2022; Zhang et al., 2018), this indicates possible cell-type specific functions of Samd4b in regulating the cilia/Shh pathway in zebrafish.

BLTP1 (KIAA1109) has been shown to have a role in endosomal trafficking and actin skeleton regulation, and its mutation causes Alkuraya-Kucinskas syndrome (Gueneau et al., 2018; Kane et al., 2019). Phenotypic manifestations of this disorder include neurodevelopmental and skeletal malformations, phenotypes commonly found in ciliopathy patients, in agreement with the defective Shh signaling following BLTP1 depletion reported here. Moreover, our data suggest a role for BLTP1 in regulation of primary cilia length. This is in agreement with observation of notably elongated cilia in fibroblasts derived from a patient harboring a *BLTP1* mutation (Kane et al., 2019). Unlike *samd4b* mutant, we did not observe any apparent gross morphology phenotypes in *kiaa1109* mutant fish. This is in contrast to skeletal malformations (including ventral curvature) reported for *bltp1* morphants (Gueneau et al., 2018), yet in agreement with no phenotype of the CRISPR-generated *bltp1* mutant reported in the same study. This indicates activation of compensatory mechanisms in *bltp1* mutant, as reported for other cilia-related mutants in zebrafish (Cardenas-Rodriguez et al., 2021; Rouf et al., 2023).

## Conclusions

Shh signaling pathway acts through primary cilia to regulate several key aspects of vertebrate development. Emerging evidence suggests that RNA-binding proteins and their regulators are found in the vicinity of cilia and the centrosome, with the functional consequence of that remaining unclear. Using proximity-based proteomics followed by functional validation, we identified SAMD4B, a previously characterized RNA-binding protein associated with cytoplasmic RNA condensates, as a centrosome-associated factor that is required for efficient Shh signaling in NIH 3T3 cells and normal eye development in zebrafish. These findings establish a previously unrecognized link between SAMD4B and ciliary signaling.

## Acknowledgements

We are grateful to Marcel Hubinský for help with custom siRNA design and to Andrea Lacigová for assistance with stable cell line generation. We thank Vladimir Varga’s lab for sharing an unpublished reagent, and James Sillibourne and Timo Glatter for CEP89/CEP123 cDNA and CP110 cDNA, respectively. We acknowledge the core facility CELLIM, supported by the Czech-BioImaging large RI project (LM2023050 funded by MEYS CR), for their support with obtaining scientific data presented in this paper. Furthermore, the CEITEC Proteomics core facility is gratefully acknowledged for obtaining the scientific data presented in this paper. CIISB, Instruct-CZ Centre of Instruct-ERIC EU consortium, funded by MEYS CR infrastructure project LM2023042 and European Regional Development Fund-Project „Innovation of Czech Infrastructure for Integrative Structural Biology” (No. CZ.02.01.01/00/23_015/0008175), is acknowledged for the financial support of the measurements at the CEITEC Proteomics core facility. Computational resources were provided by the e-INFRA CZ project (ID:90254), supported by MEYS CR.

## Contributions

OB – proximity labeling experiments, initial functional validation, assembly of figures, manuscript writing; DV – zebrafish work, manuscript writing; PP – functional validation experiments in cell lines, assembly of figures, manuscript writing; TR – functional validation experiments in cell lines; PK – functional validation experiments in cell lines; LC – project design and supervision, fund raising, manuscript writing. All authors commented on and approved the final version of the manuscript.

