## Supplementary Figures for "Identification of SAMD4B as a Centrosome-Associated Regulator of the Sonic Hedgehog Signaling Pathway"

### Supplementary Figure 1

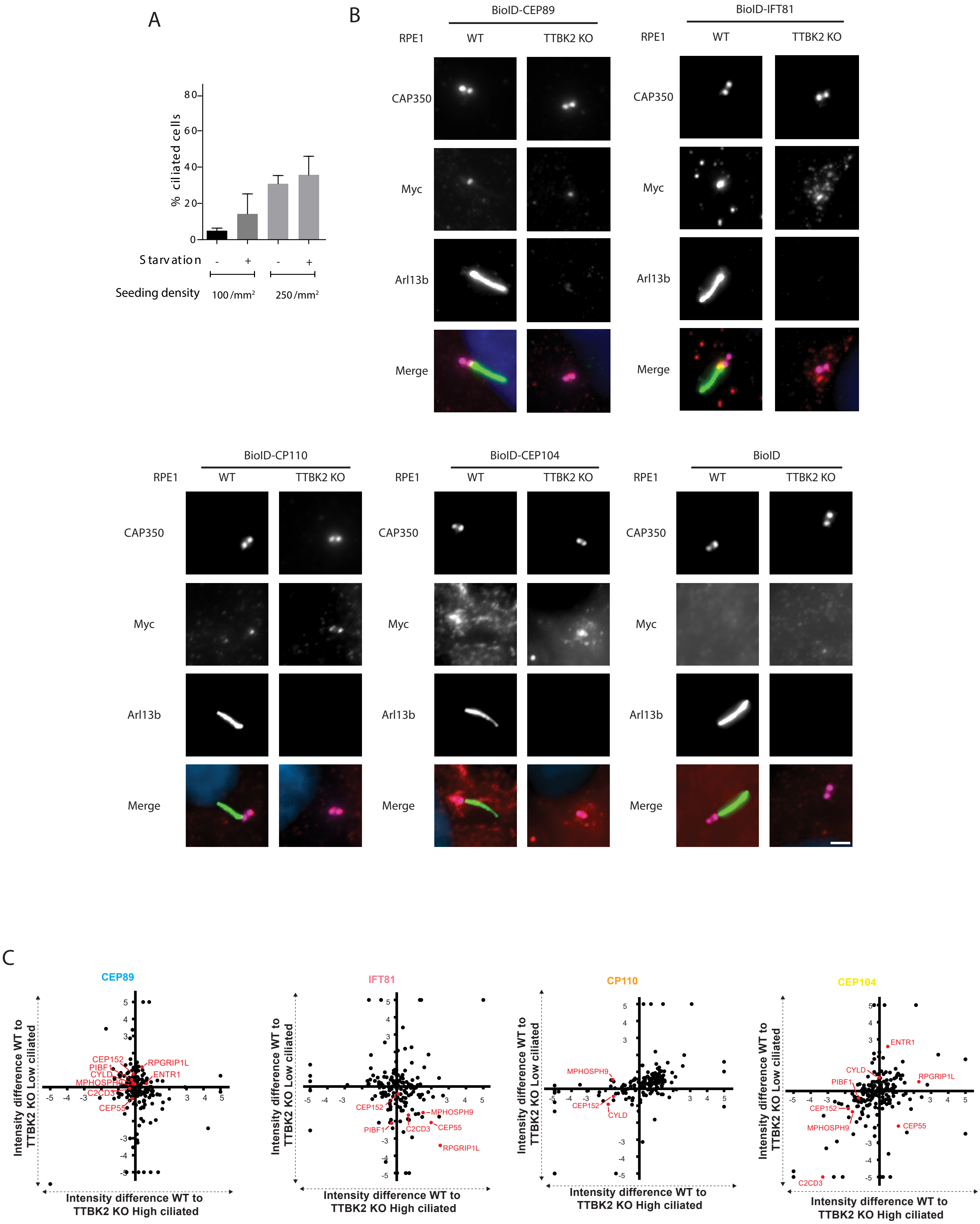

**Supplementary Fig.1:**

**A)** hTERT-RPE1 cells were seeded at the indicated densities under starved and non- starved conditions and analyzed for ARL13B-positive cilia count. The graph represents a summary of 4 independent experiments; results are shown as mean and SD.

**B)** IF validation of bait proteins N-terminally tagged with Myc-BirA (BioID) in WT and TTBK2 KO hTERT-RPE1. Cells were seeded, treated with dox, and stained for Myc (bait, in red), CAP350 (centrioles, magenta), and ARL13B (cilium, green), scale bar 1.5 µm. The BioID "only" cell line was used as a negative control.

**C)** Visualization of prey protein intensity difference between WT and TTBK2 KO. The x-axis shows the intensity difference between WT (right, positive) and TTBK2 KO (Left, negative) for the "High ciliated" dataset, while the y-axis depicts these WT (up, positive) to TTBK2 KO (down, negative) differences for the "Low ciliated" dataset. CEP89 (light blue), IFT81 (pink), CP110 (orange), or CEP104 (yellow) represent the baits used. Selected examples of detected prey proteins are highlighted in red.

### Supplementary Figure 2

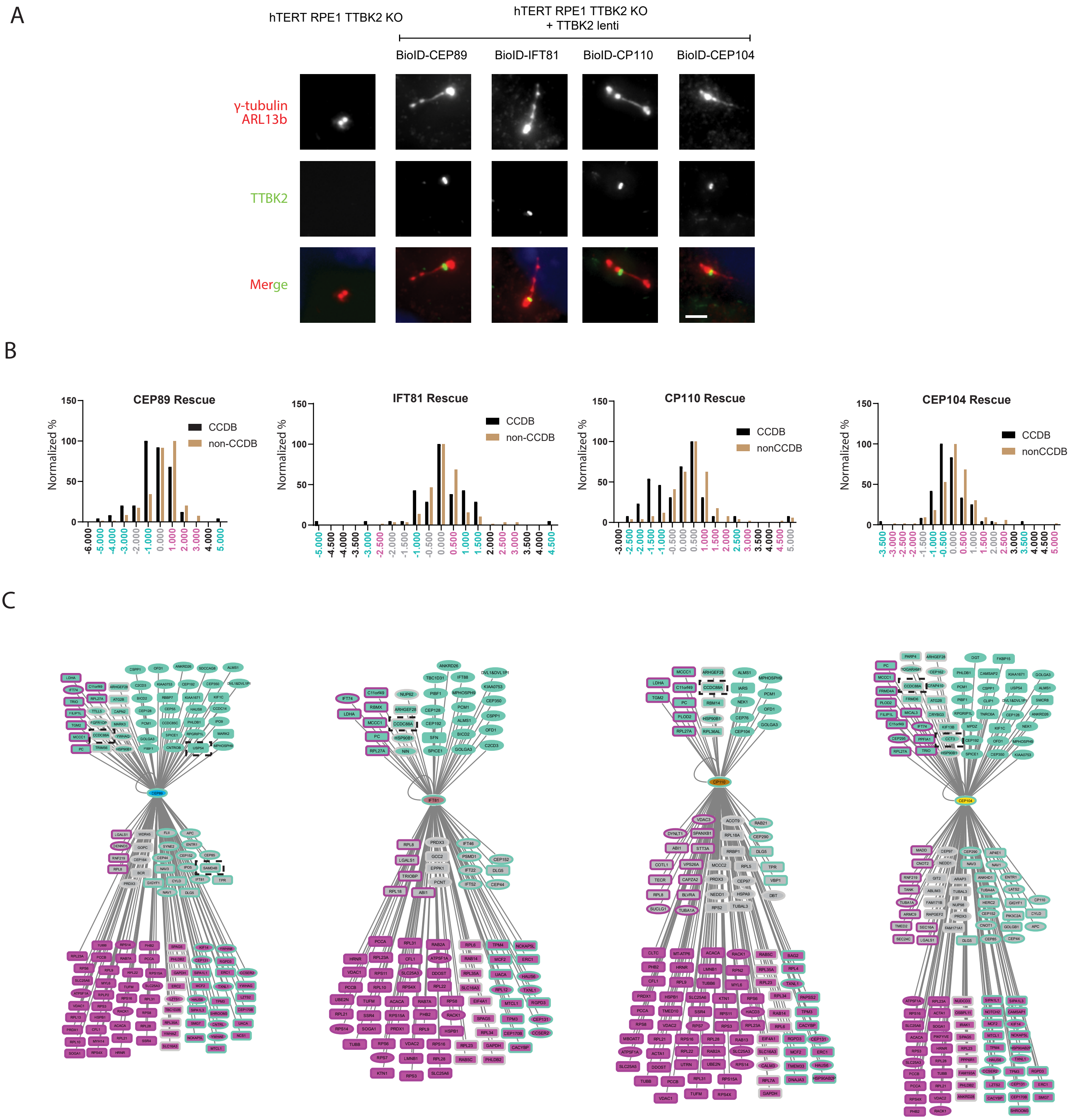

**Supplementary Fig.2:**

**A)** hTERT-RPE1 TTBK2 KO expressing indicated BioID baits were transduced by lentiviruses carrying FLAG-TTBK2 and analyzed by IF microscopy. TTBK2 reintroduction rescued ciliation of the TTBK2 KO BioID-bait cell lines and restored TTBK2 localization to the MC (green: TTBK2, red: Arl13b and γ-tubulin, blue: DAPI). Scale bar 1.5 µm.

**B)** Stratification of identified prey proteins in the TTBK2 KO rescue dataset according to their DR values. Normalized percentage is plotted on the Y axis, and the X-axis shows DW values. Prey proteins annotated as members of CCDB are indicated by black bars, whereas non-CCDB proteins are indicated by brown bars. "Enriched" groups (containing at least 30% more CCDB than non-CCDB proteins) are indicated in turquoise, "Depleted" groups (containing at least 30% fewer CCDB proteins than non-CCDB proteins) are indicated in magenta. "Neutral" groups are shown in grey.

**C)** Cytoscape-based interactomes of used bait proteins (depicted as the central node of each interacome; CEP89 - light blue, IFT81 - pink, CP110 - orange, or CEP104 - yellow) comprising both CCDB and non-CCDB proteins found across all datasets (T/DW and TTBK2 KO rescue), sorted as either "Enriched" (turquoise), "Depleted" (magenta) or "Neutral" (grey). Color of the inside of the node corresponds to the T/DW analysis, and the color of the node border reflects analyses of the TTBK2 KO rescue dataset. CCDB proteins are shown as circles, non-CCDB proteins as rectangles. Black dashed rectangles indicate proteins selected for functional analysis.

### Supplementary Figure 3

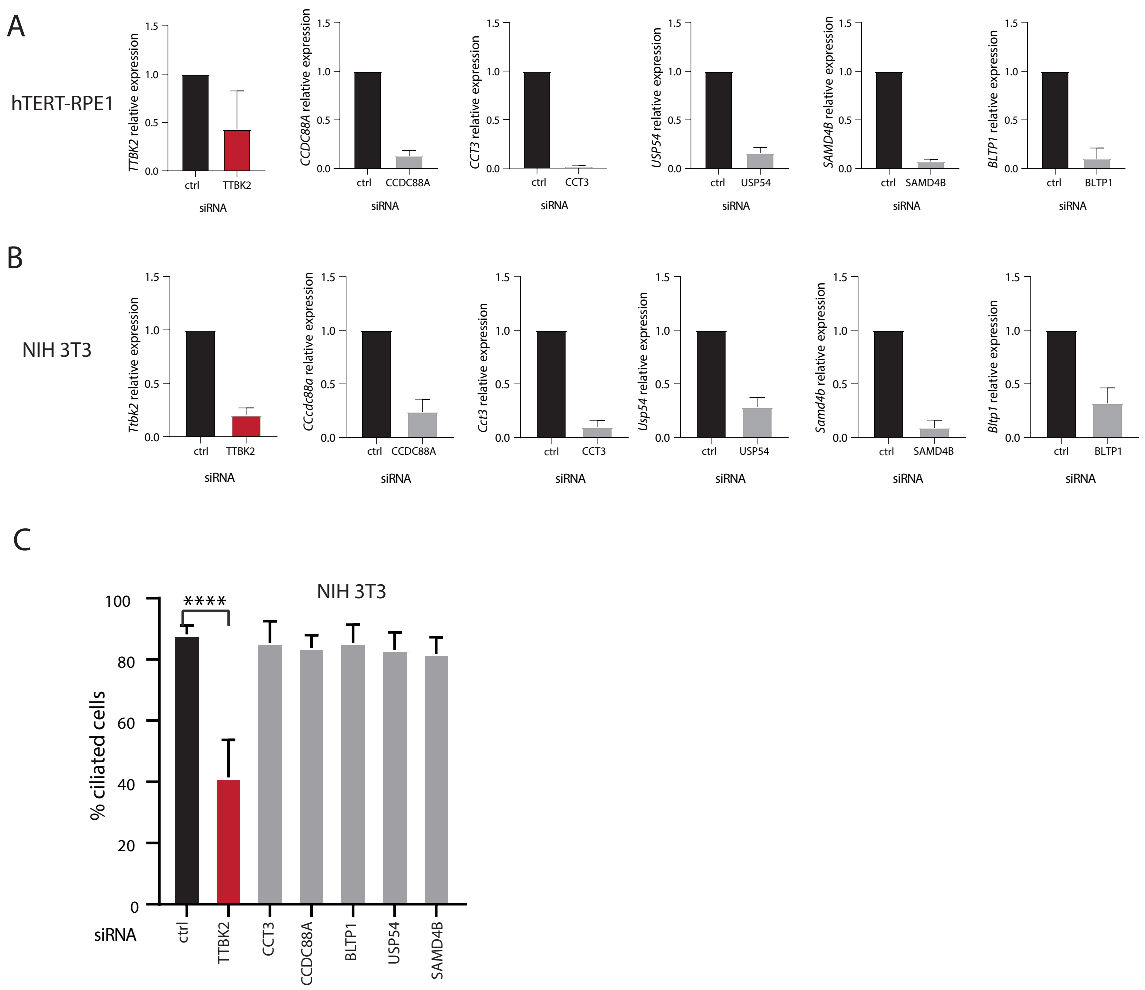

**Supplementary Fig.3:**

Q-PCR analyses of expression of indicated transcripts following siRNA transfection in **A)** hTERT-RPE1 and **B)** NIH 3T3 cells. Each graph represents a summary of at least three independent experiments. Mean and SD are shown.

**C)** Quantification of the percentage of ciliated NIH 3T3 cells in the indicated conditions. The graph represents a summary of three independent experiments, with at least 200 cells analyzed in each experiment, with a mean and SD shown. **** $p<0.0001$ by one-way ANOVA.

Supplementary Figure 4

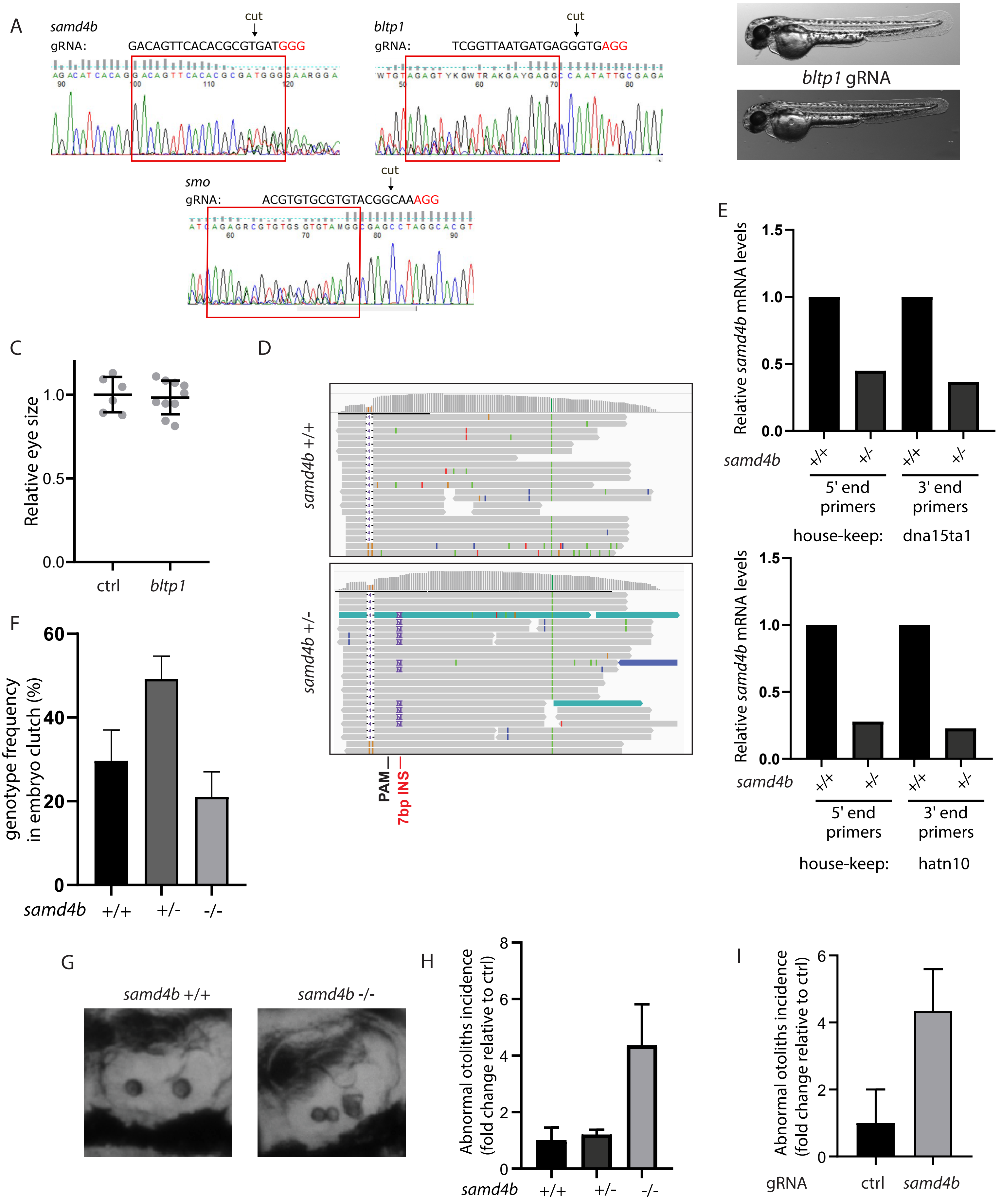

**Supplementary Fig.4:**

**A)** DNA sequencing data visualized using FinchTV, showing the ability of the used gRNAs to generate indels in the target genes.

**B)** Zebrafish embryos were injected with the indicated gRNA or mock and analyzed at about 60 hpf. Microphotographs illustrate no visible morphological differences between *bltp1* crispants and controls.

**C)** Quantification of the eye size of *bltp1* crispant fish relative to the control. Each datapoint in the graph represents one analyzed fish, with a mean and SD shown for each condition. P = 0.76 by two-tailed t-test.

**D)** NGS reads from wild-type (upper panel) and heterozygous *samd4b* 7bpINS mutant fish (lower panel) in the F1 generation (offspring of F0 founder crispants outcrossed to wild type). The 7bp insertion, which corresponds to a frameshift mutation in the *samd4b* 7bpINS allele, is marked in red and purple, respectively.

**E)** Q-PCR analyses of relative *samd4b* mRNA levels using two sets of primers and two housekeeping genes for normalizing the expression. Each condition represents a sample of 10 pooled fish from about 60 hpf.

**F)** At 60hpf, 20 embryos per clutch were randomly selected and genotyped. The graph shows the incidence of the indicated genotypes in 4 independent embryo clutches. Note that all 3 genotypes follow the expected Mendelian distribution for heterozygous incross.

**G-H)** Zebrafish larvae of the indicated genotypes were analyzed at about 60 hpf. **G)** Microphotographs illustrating abnormal otolith formation in *samd4b* -/-. Incidence of otolith abnormalities (fold change related to *samd4b* +/+) is quantified in **H)** The graph represents a summary from two independent experiments, with a mean and SEM shown. P=0.16 (+/+ versus -/-) by two tailed t-test.

**I)** Zebrafish embryos were injected with the indicated gRNA or mock and analyzed at about 60 hpf stage. The graph shows the relative incidence of otolith abnormalities (a fold change relative to ctrl), as a summary from three independent experiments, with a mean and SEM shown. P=0.12 by two-tailed t-test.

Supplementary Figure 5

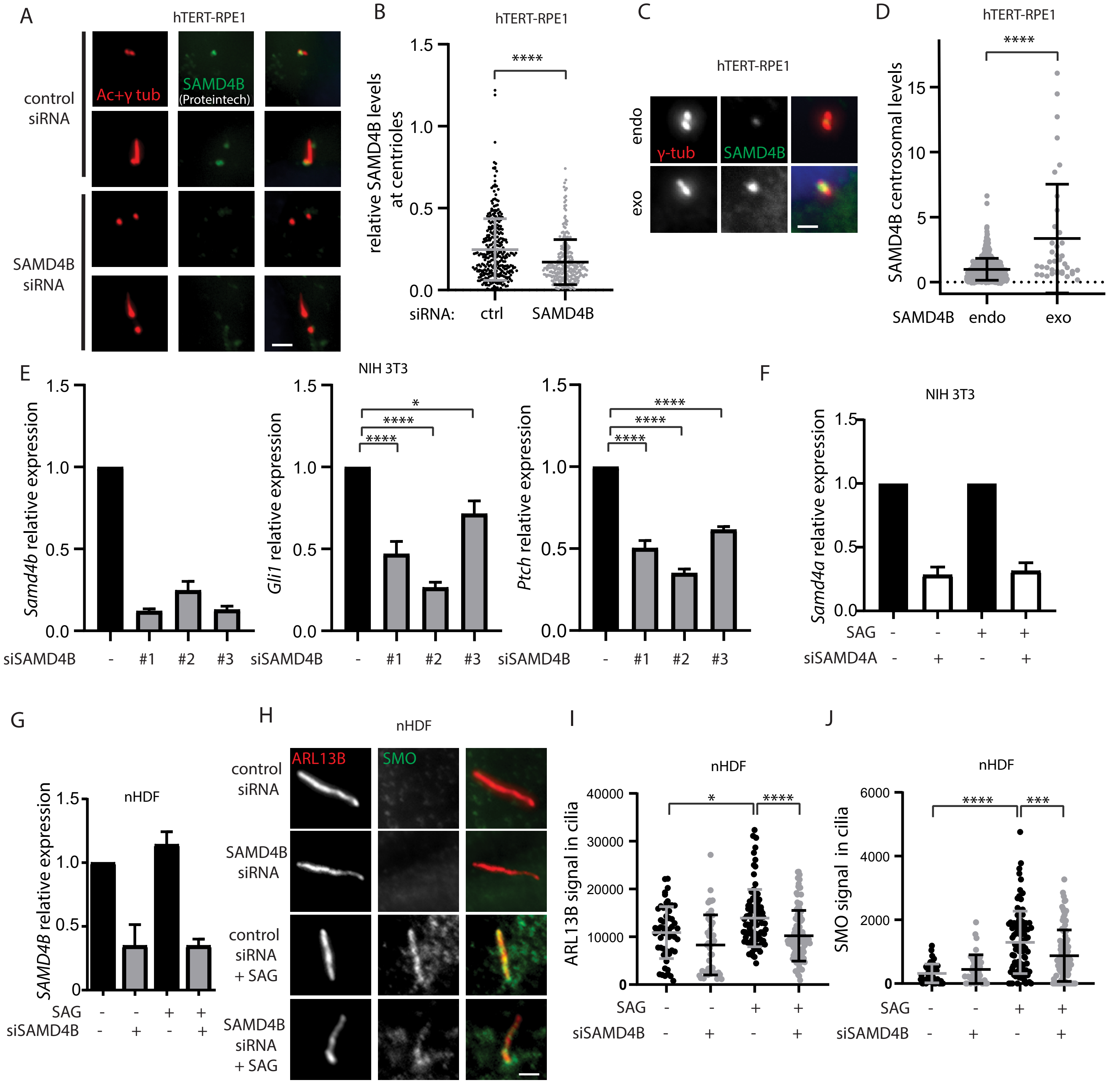

**Supplementary Fig.5:**

**A-B)** hTERT-RPE1 cells, transfected with the indicated siRNA, were fixed and stained for SAMD4B (green; Proteintech antibody) and acetylated tubulin and γ-tubulin (red) to visualize cilia and centrioles. Representative images are shown in **A),** scale bar 1.5 µm. Quantification of the relative SAMD4B signal intensity at centrioles (signal of SAMD4B normalized to signal of acetylated tubulin and γ-tubulin) is shown in **B).** The graph represents a summary of five independent experiments, with individual datapoints, mean, and SD shown. **** $p<0.0001$ by two-tailed t-test.

**C-D)** hTERT-RPE1 cells were transfected with SAMD4B plasmid, fixed, and stained for SAMD4B (green) and γ-tubulin (red). Representative images are shown in **C)**, scale bar 1 µm. Note that transfected cells showed higher levels of SAMD4B associated with centrioles (exo) than non-transfected controls (endo). Quantification of the SAMD4B signal intensity is shown in **D).** The graph represents a summary of three independent experiments, with individual datapoints, mean, and SD shown. $p<0.0001$ by two-tailed t-test.

**E-G)** Q-PCR analyses of relative expression of indicated transcripts following siRNA transfection, treated with 200nM (NIH 3T3) or 500 nM (nHDF) SAG, where indicated. **E-F)** Each graph represents a summary of at least three independent experiments in NIH 3T3, with mean and SEM shown. **** $p<0.0001$, **$p<0.01$, *$p<0.05$ by one-way ANOVA. **G)** The graph represents a summary of two independent experiments in nHDF, with mean and SEM shown.

**H-J)** nHDF cells, transfected with the indicated siRNA and treated with vehicle or 500 nM SAG, were fixed and stained for SMO (green) and ARL13B (red). Representative images are shown in **H),** scale bar 1.5 µm. Quantification of the ARL13B signal intensity in cilia is shown in **I)**. Quantification of the SMO signal intensity in cilia is shown in **J)**. Each graph represents a summary of two independent experiments, with individual datapoints, mean, and SD shown. **** $p<0.0001$, *** $p<0.001$, * $p<0.05$ by one-way ANOVA.
